# Hotspot *KRAS* mutations in brain metastases at the first metastatic recurrence of cutaneous melanoma

**DOI:** 10.1101/2020.02.17.952630

**Authors:** Roy Rabbie, Peter Ferguson, Kim Wong, Una Moran, Clinton Turner, Patrick Emanuel, Kerstin Haas, Jodi M. Saunus, Morgan R. Davidson, Sunil R. Lakhani, Brindha Shivalingam, Georgina V. Long, Christine Parkinson, Iman Osman, Richard A. Scolyer, Pippa Corrie, David J. Adams

**Author notes:** **Correspondence to**: Dr. David Adams, Experimental Cancer Genetics, Wellcome Sanger Institute, Hinxton, Cambridge, CB10 1HH, **Ph**, **Email**. These authors contributed equally to this work (Roy Rabbie, Peter Ferguson).

## Abstract

**IMPORTANCE:** Brain metastases occur in 60% of patients with advanced melanoma and are a major cause of melanoma-related mortality and morbidity. Although our understanding of the molecular alterations associated with melanoma progression is improving, there are currently no validated biomarkers which might help identify those patients at highest risk of developing brain metastases.

**OBJECTIVE:** To examine the somatic mutational and copy-number landscape of brain metastases that develop as the isolated first visceral site of recurrence – “early brain-metastasis” compared to extracranial melanoma metastases.

**DESIGN, SETTING AND PARTICIPANTS:** Whole-exome sequencing of 50 tumors from patients undergoing surgical resection of one or more brain metastasis occurring as the first site of visceral relapse were identified from prospectively maintained databases in Sydney, Wellington, New York and Cambridge. Whole exome sequencing analyses allowed mutational profiles to be compared to cutaneous melanomas in The Cancer Genome Atlas (SKCM-TCGA; n=358) and the Memorial Sloan Kettering (SKCM-MSK-IMPACT; n=186) datasets. An external dataset comprising a further 18 patients with surgically resected early brain metastasis from two additional academic centers served as an independent validation cohort.

**MAIN OUTCOMES AND MEASURES:** To assess the frequency of driver mutations in early brain metastasis and their influence on survival.

**RESULTS:** In concordance with the landmark melanoma sequencing studies, we identified mutations in BRAF (21/50, 42%), NRAS (14/50, 28%) and NF1 (11/50, 22%) as the most frequently mutated melanoma driver genes. When compared to the mutational landscape of cutaneous melanomas in TCGA (SKCM-TCGA), KRAS was the most significantly enriched driver gene, with 5/50 (10%) of brain metastases harboring non-synonymous mutations, of which 4/5 (80%) were in the hotspot positions of codons 12 and 61. This was significantly higher than the corresponding frequency of *KRAS*-mutations within the entire SKCM-TCGA (2% (7/358), p=0.009, Fisher’s Exact Test) as well as the SKCM-MSK-IMPACT cohort (1.6% (3/186), p=0.016). Variants in KRAS were mutually exclusive from *BRAF*^*V600*^, *NRAS* and *HRAS* mutations and were associated with a significantly reduced overall survival from resection of brain metastasis (relative to *KRAS*-wild type brain metastases) in multivariate Cox proportional hazard models (HR 1.80, 95% CI 1.46-24.89, p=0.013). Mutations in *KRAS* were also clonal and concordant with extracranial disease, which suggests these mutations are present within the primary tumor

**CONCLUSIONS AND RELEVANCE:** Our analysis, the largest to date, suggests that early metastases to the brain (presenting as the first site of visceral relapse) are characterized by significant enrichment of hotspot *KRAS* mutations, potentially implicating constitutive RAS-driven cellular programs in neurotropic metastatic behavior in these cases. Based on these data, we suggest that screening for *KRAS* mutations might help identify those patients with primary melanoma at higher risk of brain metastases or poor survival, and could help inform future surveillance strategies.

**Key Points:** *Question:* What is the frequency of driver mutations in early melanoma brain metastases?

*Findings:* In this study of 50 patients with melanoma metastasizing first to the brain, *KRAS* mutations were the most significantly enriched driver gene (n=5, 10% of patients) when compared to landmark cutaneous melanoma studies. The high *KRAS* mutation frequency was also observed in an external validation cohort of 18 patients with early brain metastases. Mutations in *KRAS* were mutually exclusive from mutations in the key RAS signaling genes and conferred a worse overall survival from resection of brain metastasis.

*Meaning:* Hotspot *KRAS* mutations could help identify those patients with primary melanoma at higher risk of brain metastases that may benefit from more intensive, protracted surveillance as well as earlier use of adjuvant therapy.

## Background

Metastases to the central nervous system (CNS) are observed in ~60% of cutaneous melanoma patients developing disseminated disease and up to 90% at autopsy.^1^ Early detection of intracerebral recurrences remains critical, as isolated or oligometastatic brain metastases may be more amenable to potentially curative locoregional therapies and immunotherapies have demonstrated greatest efficacy in patients with small, asymptomatic metastases.^1–3^ Early predictors of brain metastases could therefore help identify those patients most likely to benefit from closer surveillance of the brain as well as early use of adjuvant therapies.

Importantly, epidemiological data suggest that patterns of metastatic dissemination may be partially determined by the clinical characteristics of the primary tumor (e.g. scalp/high mitotic rate primary melanomas might have higher risk of brain metastases^4^). However, melanomas have considerable molecular diversity and several lines of evidence suggest that metastatic melanomas (including brain metastases) are genetically evolved from their primary tumor predecessors.^5^

Interestingly, 15-20% of brain metastases present as the isolated first visceral site of recurrence.^6^ Primary tumors in these ‘early brain metastasis’ cases reported as thinner and of lower American Joint Committee of Cancer (AJCC) Stage when compared to other visceral metastases, challenging the current understanding of brain metastases as the final stage of tumor progression and suggesting these tumors could harbor distinct biological properties favoring early hematogenous dissemination to the brain.^6^ Our analyses of the mutational landscape of early brain metastasis highlights key molecular features that could inform future prognostic, surveillance and intervention strategies.

## Methods

### Study population

Patients with available archival paraffin embedded melanoma brain metastases (in the absence of other sites of visceral disease, confirmed by CT or MRI imaging prior to neurosurgery) were selected from prospectively maintained databases at The Melanoma Institute of Australia (n=34), The Wellington School of Medicine (n=8), New York University School of Medicine (n=4) and Cambridge University Hospitals (n=4) (discovery cohort). Samples from patients selected from The University of Auckland and The University of Queensland Australia (n=18 total) made up the external validation cohort. All neuro-resections were undertaken between 2008 and 2018 at the respective academic neurosurgical centers as part of routine clinical care. All cases were ethically approved by the local Institutional Review Boards, as well as by the Sanger Institute’s human materials and data management committee. All samples and clinical details are listed in **Supplementary Table 1**.

The clinical and mutation data from The Cancer Genome Atlas (SKCM-TCGA)^7^, was downloaded from the cBioPortal. The SKCM-MSK-IMPACT dataset was extracted from the publication by Zehir *et al*^8^ (**Supplementary Methods**).

### DNA sequencing

Exome capture of the discovery cohort was performed using Agilent SureSelect All Exon V5 bait. Paired-end sequencing was performed using the Illumina HiSeq (Illumina, San Diego, CA, USA) platform at the Wellcome Sanger Institute. In (n=16) cases where no matching germline DNA was available, a panel of 39 FFPE-extracted normals were used to filter germline variants as well as artefacts. MuTect (v1.1.7) and Sequenza (v2.1.2) were used to call somatic SNVs and copy number aberrations, respectively. Panel sequencing of the 18 external validation cohort samples was performed using custom pull-down sequencing (**Supplementary Methods**).

### Survival analyses

Univariate Kaplan-Meier survival plots were constructed using the R ‘Survival’ package version 2.4.2. Univariate and multivariate hazard ratios (HR), 95% confidence intervals (95% CI) and corresponding p-values were obtained using Cox proportional hazards regression models. Multivariate corrections were undertaken for gender, age, as well as BRAF and NRAS mutation status.

## Results

Fifty patients who developed brain metastases as their first site of visceral disease spread were enrolled as part of the discovery cohort and were represented by a relatively high proportion of thin (T1-T2) (n=25, 50%) and non-ulcerated (n=26, 52%) primary melanomas (**Supplementary Table 1**).

Mutations in BRAF were detected in 21 (42%) tumors, of which 15/21 (71%) were in the V600 hotspot (**Figures 1A & 1B**). NRAS mutations were identified in 14 (28%) tumors and were all in hotspot positions on exons 2 (codons 12 and 13) and 3 (codon 61) and mutually exclusive from *BRAF*^V600^ mutations. Comparing the mutational landscape of brain metastases to that of cutaneous melanomas in the SKCM-TCGA dataset (see **Supplementary Methods**), *KRAS* was the most significantly enrichment driver gene in our dataset, mutated in 10% (5/50) vs 2% (7/358) in the entire SCKM-TCGA collection, p=0.009 two-tailed Fisher’s exact test). The mutation frequency of KRAS was also significantly enriched relative to the frequency of extracranial melanoma metastases, 10% (5/50) in our dataset vs 2.1% (6/274) in extracranial melanoma metastases in SKCM-TCGA (p=0.016, two-tailed Fisher’s exact test, see **Supplementary Methods**). Further, only 1.6% (3/186) of melanoma cases in the SKCM-MSK-IMPACT dataset^8^ were *KRAS* mutant, significantly lower than in our early brain metastasis discovery cohort (p=0.016 two-tailed Fisher’s exact test). Mutations in *KRAS* had a high variant allele frequency, indicating that these likely represent clonal (early) driver mutations (**Table 1**). Of note, three extracranial metastases available for sequencing from patients with *KRAS*-mutant brain metastases also harbored the same brain-metastatic *KRAS* mutations, suggesting that KRAS mutations were concordant with extracranial metastases (see **Supplementary Methods**). Notably, four out of five of the brain metastases mutations in *KRAS* were in hotspot codons 12 and 61 and mutually exclusive from other mutations in the mitogen-activated protein kinase (MAPK) signaling genes including *BRAF*^*V600*^, *NRAS* and *HRAS* (**Figure 1B**).

**Table 1.** Clinical and mutational characteristics of the patients with hotspot *KRAS* mutations (including both patients from the discovery and validation cohorts). Of note all hotspot *KRAS* mutations were identified in patients with either thin (T1/T2) or no prior history of primary melanoma and from primary tumors in chronically sun-exposed locations. Mutations in *KRAS* also had a high variant allele frequency, indicating that these likely represent clonal driver mutations. Cambridge: Cambridge University Hospitals, MIA: Melanoma Institute of Australia, Queensland: The University of Queensland Australia. SSM: Superficial spreading melanoma, NM: Nodular melanoma, LMM: Lentigo-maligna melanoma.

| Patient_Pin | SampleID | Centre | Gender | Primary site | Histologic subtype | Breslow thickness (mm) | Ulceration | Clark level | Mitoses | Tstage | KRAS mutated AA position | Variant allele frequency |
| --- | --- | --- | --- | --- | --- | --- | --- | --- | --- | --- | --- | --- |
| MBM_Disc_10 | PD42091a | MIA | Female | Extremity | SSM | 0.2 | No | II | 0 | T1 | G12S | 0.85 |
| MBM_Disc_23 | PD42097a | MIA | Male | Head/neck | NM | 1.5 | No | NA | NA | T2a | Q61R | 0.68 |
| MBM_Disc_27 | PD42100a | MIA | Male | Head/neck | LMM | 1.5 | Yes | IV | 5 | T2a | Q61L | 0.86 |
| MBM_Disc_40 | PD31211c | Cambridge | Male | Head/neck | Melanoma-in-situ | 0 | No | I | 0 | T1a | G12D | 0.50 |
| MBM_Valid_10 | PD44959a | University of Queensland | Male | No history primary | No history primary | No history primary | No history primary | No history primary | No history primary | No history primary | G13C | 0.70 |

**Figure 1.**
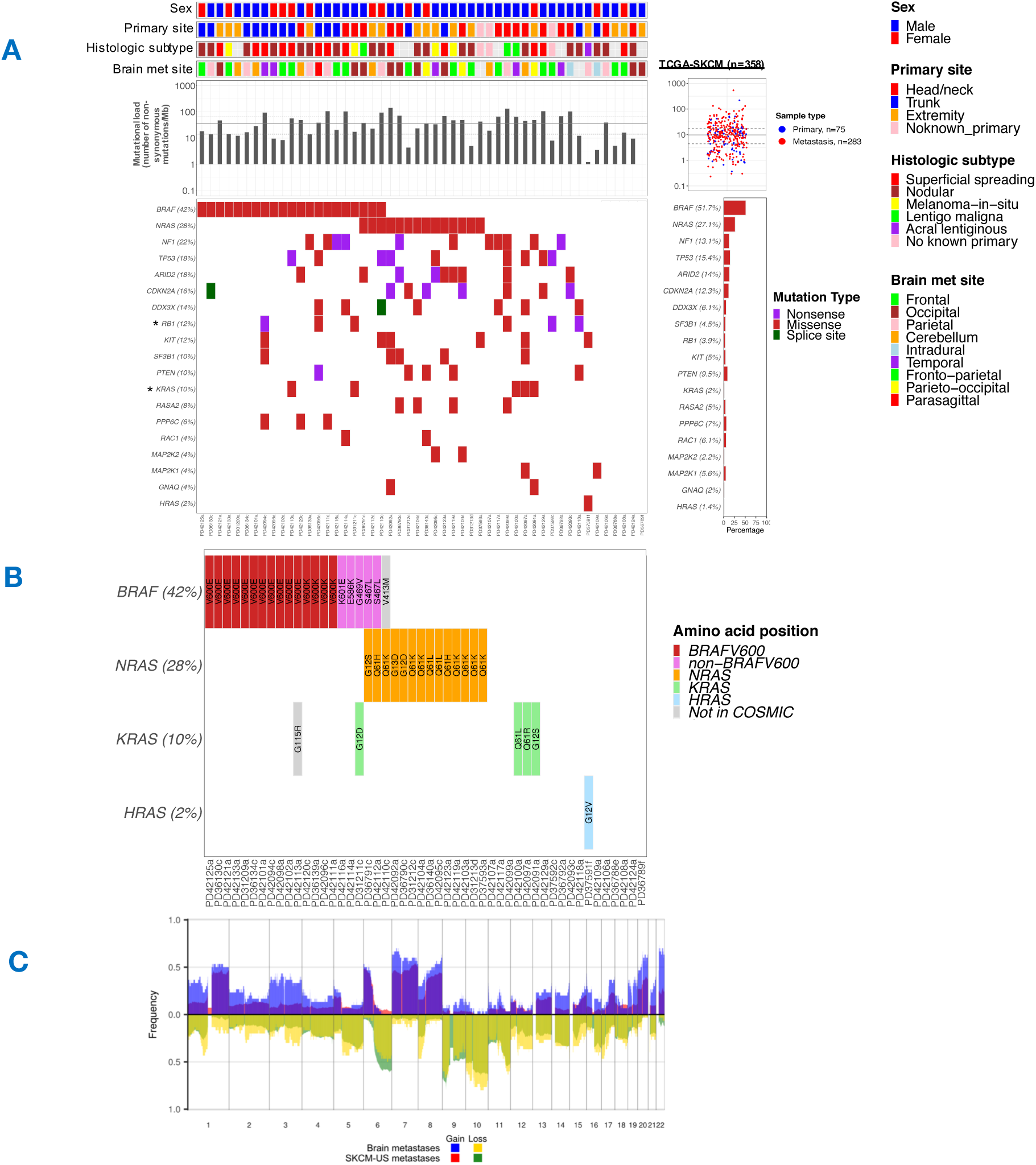
Tile plot of melanoma driver mutations in the early brain metastasis discovery cohort (n=50). **A)** The mutational profiles of brain metastases are indicated. Mutational load was calculated as the number of non-synonymous mutations per Mb, median is indicated by the solid horizontal grey line and the 95% confidence interval by the dashed lines (median 38 mutations/Mb, 95% CI 14.2-67.1). The genes shown carry non-synonymous mutations within the selected melanoma drivers outlined in Hayward *et al*. (n=19) and are ordered according to their mutation frequency within this cohort. The corresponding mutational load and gene mutational frequencies in the SKCM-TCGA dataset (n=358) is indicated in the plot aside. SKCM-TCGA mutational load boxplots are colored according to whether the sample was classified as a primary (n= 75, blue) or metastasis (n=283, red). The median non-synonymous mutational load in SKCM-TCGA is indicated by the solid horizontal grey line and the 95% confidence interval by the dashed lines (median 9.9 mutations/Mb, 95% CI 4.4-18.3). *signifies the gene is significantly more mutated in the early brain metastasis cohort. *KRAS*; 10% (5/50) vs 2% (7/358) in the SCKM-TCGA collection, p=0.009 Fisher’s exact test), *RB1*; (6/50 12% in our dataset vs 3.9% 14/358 in SKCM-TCGA, p=0.025 Fisher’s exact test). **B)** Focused tile plot from (A), highlighting the mutated amino acid positions within the *RAS* signaling genes. As expected, mutations in *NRAS* were mutually exclusive to *BRAF*^*V*600^ hotspots. The four hotspot *KRAS* mutations were also mutually exclusive to *BRAF*^V600^ and to mutations in *NRAS* and *HRAS*. Mutations shaded in grey were not found in the cBio catalogue of cancer mutations and likely represent passenger mutations. **C)** Copy number profile of the early melanoma metastasis discovery cohort (n=30) overlaid onto the copy number profile of SKCM-TCGA (n=337). The non-overlaid plots are shown in **Supplementary Figure 2.**

We conducted a further custom pull-down validation experiment on selected melanoma driver mutations within the discovery cohort and confirmed 56/60 (93%) to be somatic mutations (see **Methods**). We also conducted another external validation experiment, analyzing a further 18 early metastases independently acquired from two different neurosurgical centers (see **Supplementary Methods**). This revealed 1 brain metastasis (5.6%) harbored a *KRAS*^*G13C*^ mutation, which was also mutually exclusive from mutations in the RAS signaling genes (*BRAF/NRAS/HRAS*) (**Supplementary Figure 1**). The copy number landscape of the early brain metastasis discovery cohort proved remarkably similar to that of SKCM-TCGA (**Figure 1C & Supplementary Figure 2**).

All patients with *KRAS*-mutant brain metastases succumbed to disease, with a median overall survival from resection of brain metastasis of only 3 months, compared to 12 months in patients with resected *KRAS*-wild type brain metastases (HR 1.80, 95% CI 1.46-24.89, p=0.013, covariate corrected Cox proportional hazards model, **Figures 2A & B**). Melanoma patients with tumors harboring *KRAS* mutations or amplifications represented in the SCKM-TCGA dataset were also associated with worse overall survival compared to *KRAS* wild-type melanomas (HR 2.6, 95% CI 1.2-5.4, p=0.015 univariate Cox-regression), although this did not meet the threshold for statistical significance after correction of clinical covariates, likely due to the limited sample size (HR 2.1, 95% CI 0.9-4.7, p=0.07, multivariate corrected Cox proportional hazards regression, **Supplementary Figure 4**).

**Figure 2.**
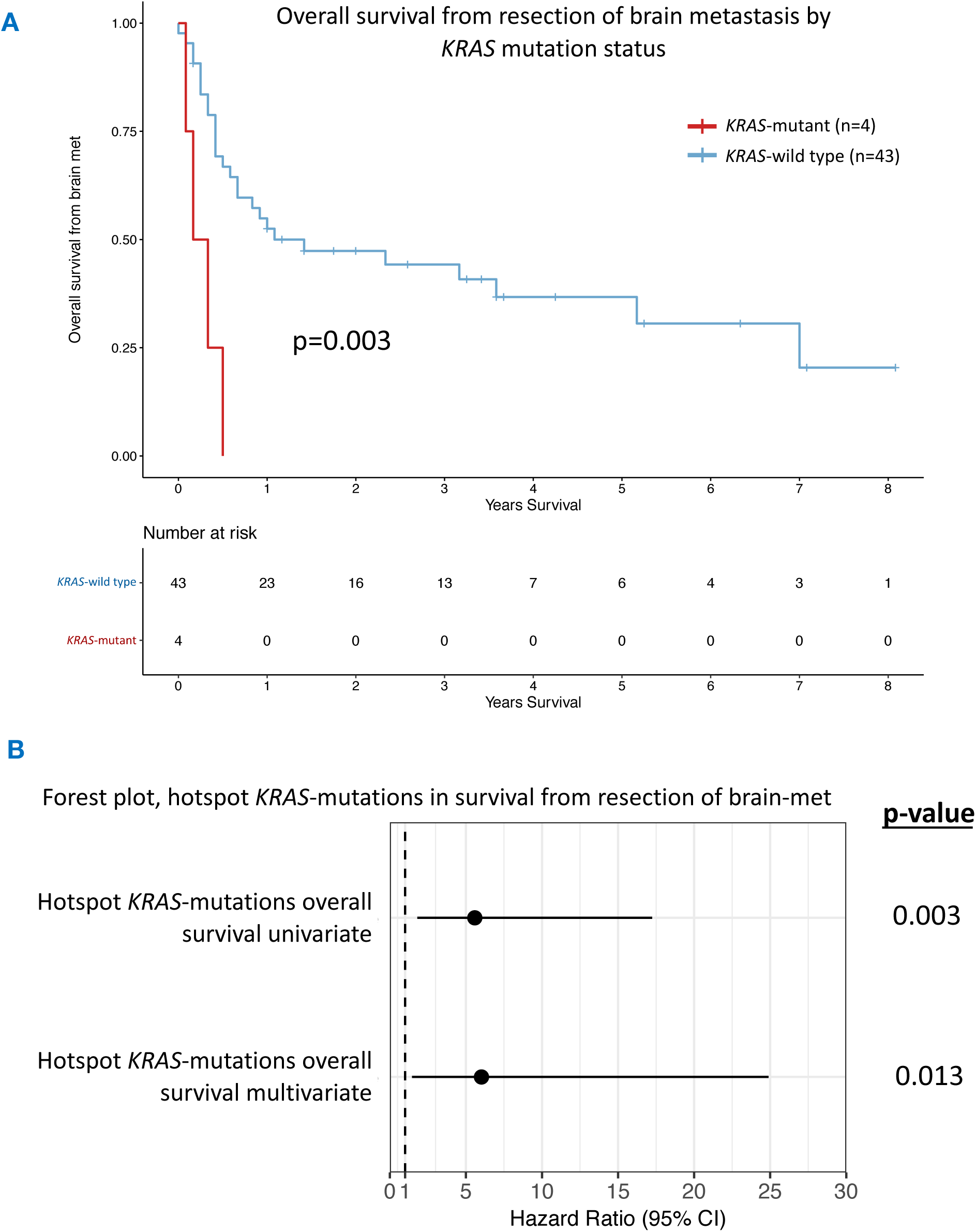
Impact of *KRAS* mutational status on survival. **A)** Kaplan-Meier survival curves showing overall survival probabilities from resection of brain metastasis (defined as the time from the resection of the brain metastasis to last follow-up (censored) or death from any cause) for the hotspot *KRAS*-mutant (n=4) versus *KRAS*-wild-type (n=43) patients (7 *KRAS* wild-type patients did not have survival data available). Patients with *KRAS*-mutant tumors had significantly worse overall survival from resection of brain metastasis than *KRAS*-wild-type patients, median 3 vs 12 months (p=0.002, univariate Cox regression). **B)** Forest plot comparing *KRAS*-mutant versus wild-type survival from resection of brain metastasis in univariate (HR 5.58, 95% CI 1.80-17.24, p=0.003) and multivariate (HR 6.02, 95% CI 1.46-24.89, p=0.013) Cox proportional hazards regression models. Multivariate correction was undertaken for gender, age at resection of brain metastasis, as well as *BRAF* and *NRAS* mutation status.

## Discussion

This analysis represents the largest survey of mutation profiles of melanoma brain metastases and the first to show an association between *KRAS* mutations and adverse outcomes in melanoma. The predominance of *KRAS* mutations in codons 12, 13 and 61, as well as the mutual exclusivity to other key drivers of MAPK signaling suggests these likely represent important drivers in this context.

The RAS family of GTPases consists of genes including *NRAS, KRAS* and *HRAS*, mutated in 25%, 2% and 1% of melanomas respectively.^7^ *NRAS*-mutant melanomas are recognized to be more aggressive and associated with poorer outcomes, however very little is known about *KRAS*-mutant melanoma.^9^ *KRAS*-mutant early brain metastasis in our study generally emanated from thin and non-ulcerated primary melanomas (**Table 1**). Hence, *KRAS* detection might in future be used to ‘upstage’ a subgroup of lower risk patients not currently offered routine surveillance and/or adjuvant therapy potentially avoiding the devastating impact of brain metastases. Mutations in *KRAS* were clonal and concordant with extracranial disease, which suggests these mutations are present within the primary tumor, however further studies will be required to confirm this.

The MAPK and phosphoinositide-3 kinase (PI3K) pathways are the two key downstream signaling pathways through which constitutively activated RAS exerts its pro-tumorigenic effects. MAPK pathway activation and brain metastases are inextricably connected and *BRAF* and *NRAS* mutations are associated with an increased risk of brain metastasis.^10^ In the same way the PI3K/AKT pathway has been mechanistically linked with the development of brain metastases^11^, and analyses of patient-matched pairs of brain and extracranial metastases have revealed that brain metastases have higher levels of activated AKT and lower expression of *PTEN*, a finding also observed using immunohistochemistry.^12^ Hotpot *KRAS* mutations are known to activate *EGFR* signaling pathways, which in-turn is associated with an increased risk of brain metastases in non-small cell lung cancer (NSCLC)^13^. One human sequencing study has further shown a direct association between *KRAS*-mutations and NSCLC brain metastases^14^, although further functional validation will be required.

## Limitations

The retrospective nature of this analysis could feasibly introduce a degree of selection bias, in particular by only identifying those patients with operable early brain metastasis we might have excluded a larger patient demographic with more widespread disease. Emerging evidence indicates that metastatic outgrowth may also depend on the interplay between cancer cells and the host stroma^15^ however such tumor-cell extrinsic factors might not be fully captured by this analysis. The identification of *KRAS* mutations as a predictive biomarker for the development of early brain metastases will ultimately require prospective validation in larger cohorts employing multivariate models, particularly assessing its predictive value in relation to other clinical covariates^4^.

## Conclusion

In summary, our analyses indicate that the patterns of melanoma recurrence may be at least partially determined by the tumor mutational profile, and that up to 10% of patients developing early brain metastases may have tumors driven by oncogenic *KRAS* mutations. This observation has implications for deciphering the biology of site-specific metastatic pathogenesis and, if validated in larger prospectively curated cohorts, might influence prognosis, surveillance and intervention in patients carrying these somatic alterations.

## Supporting information

All tables

## Acknowledgements

First and foremost, we would like to sincerely thank the patients involved in this study. We would also like to thank the Cancer, Ageing and Somatic Mutation Program team at the Sanger Institute for facilitating QC and DNA sequencing, particularly Claire Hardy, Stephen Gamble and Elizabeth Anderson. We would also like to thank Ingrid Ferreira for performing the histopathological analyses for the University of Queensland cases. Sofia Chen, James Gilbert and Matthew Garnett for their help with the custom capture bait. Assistance from colleagues at Melanoma Institute Australia and Royal Prince Alfred Hospital is gratefully appreciated. Carla Daniela Robles Espinoza for critical review of the data and the manuscript.

## Figure legends

**Supplementary Figure 1.**
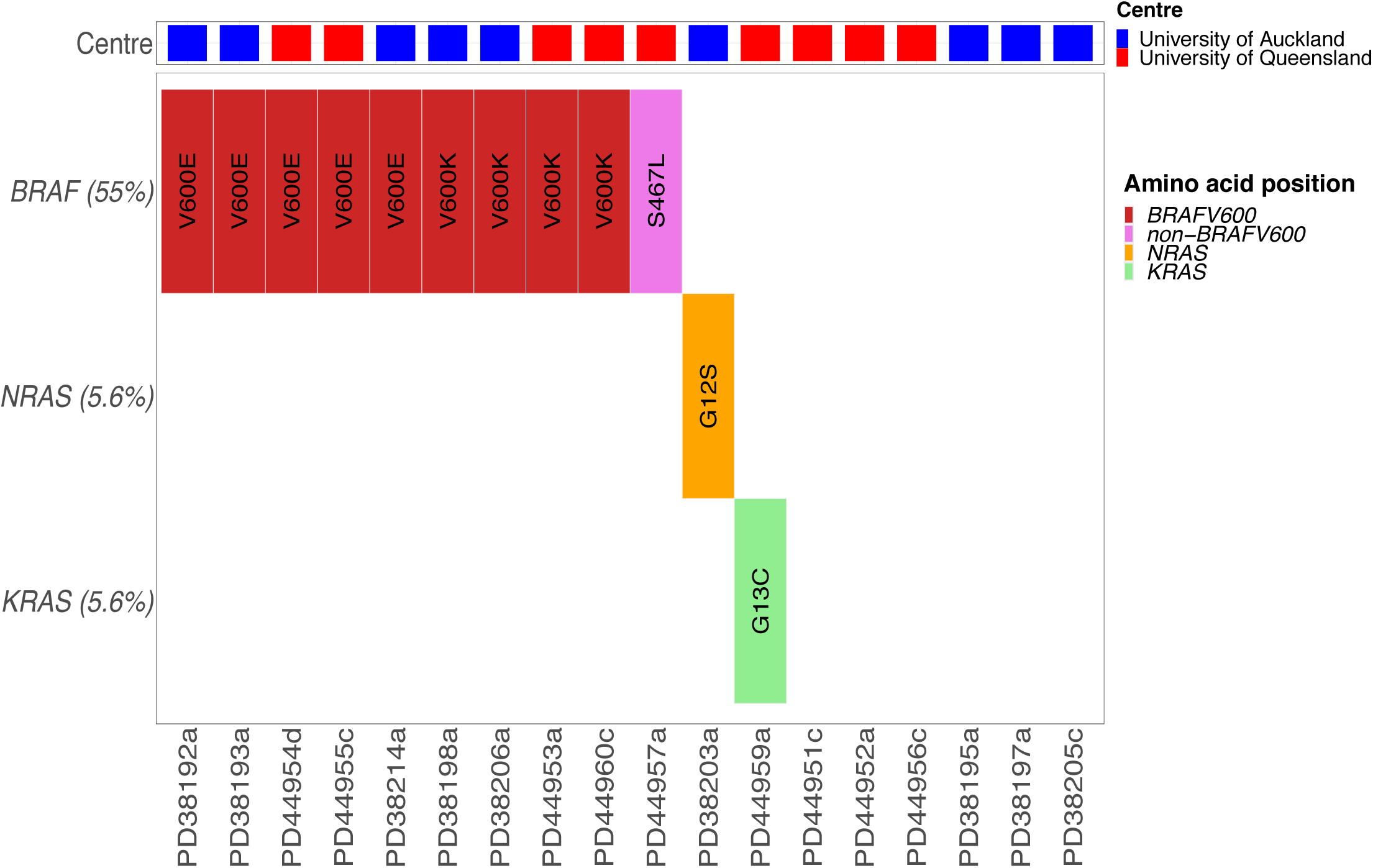
Tile plot of mutated positions within the RAS signaling genes in the external validation cohort (n=18). The *KRAS*^*G13C*^ mutation was mutually exclusive to both *BRAF* and *NRAS* hotspot mutations, in keeping with the findings from the discovery cohort (Figure 1B).

**Supplementary Figure 2.**
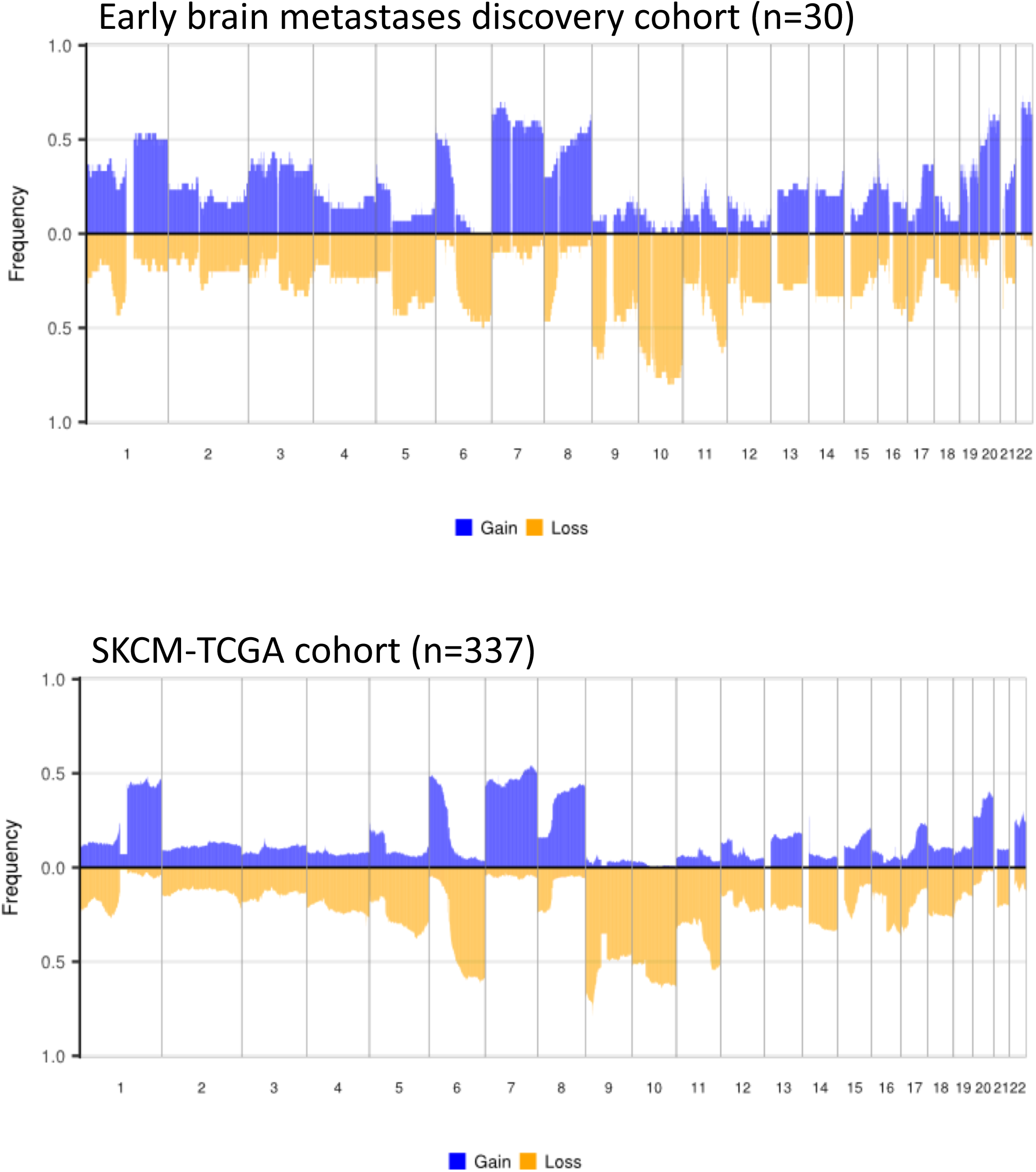
Copy number profile of early melanoma brain metastasis from the discovery cohort (top, n=30) and the SKCM-TCGA (bottom, n=337).

**Supplementary Figure 3.**
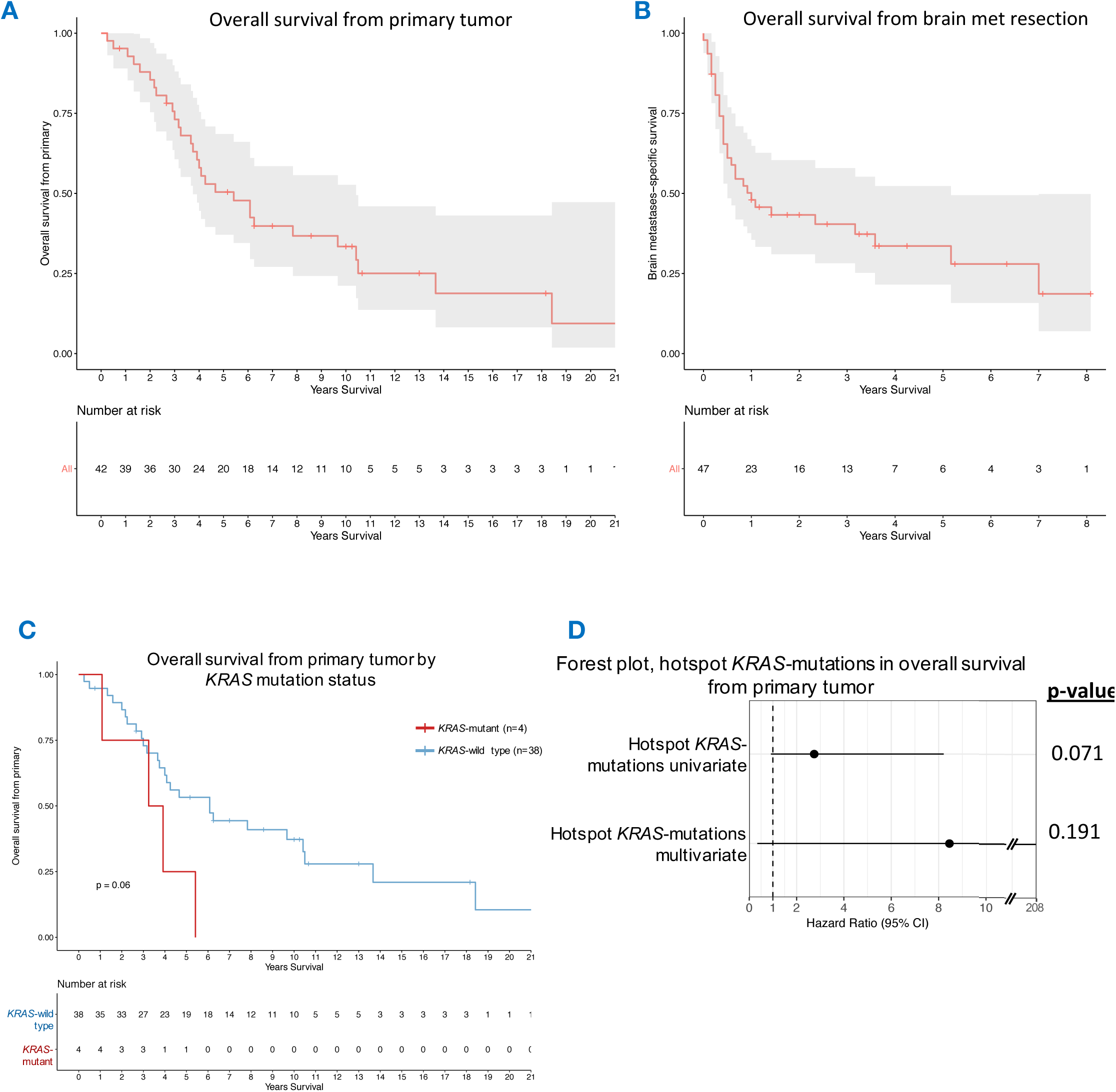
Kaplan-Meier survival curves of the early brain metastasis discovery cohort. **A**) Overall survival from primary disease (defined as the time from the resection of the primary tumor to last follow-up (censored) or death from any cause). Shaded regions represent the 95% confidence interval (median survival from primary disease 65.0 months, 95% CI 45.0-125.0 months). Survival data from primary disease was only available on 42 patients (3 patients had no history of primary melanoma, 5 patients had no survival information). **B)** Overall survival from resection of brain metastasis, defined as the time from the resection of brain metastasis to last follow-up (censored) or death from any cause. Median 12.0 months, 95% CI 6.0-62.0 months. Data from resection of brain metastasis was available on 47 patients. **C)** Impact of *KRAS* mutational status on overall survival from primary tumor. **D)** Forest plot comparing *KRAS*-mutant versus *KRAS*-wild-type overall survival from primary tumor in univariate (HR 2.74, 95% CI 0.92-8.21, p=0.071) and multivariate (HR 8.46, 95% CI 0.34-207.61, p=0.191) Cox proportional hazards regression models. Multivariate correction was undertaken for gender, age at primary tumor resection, T-stage and ulceration of primary tumor, as well as *BRAF* and *NRAS* mutation status.

**Supplementary Figure 4.**
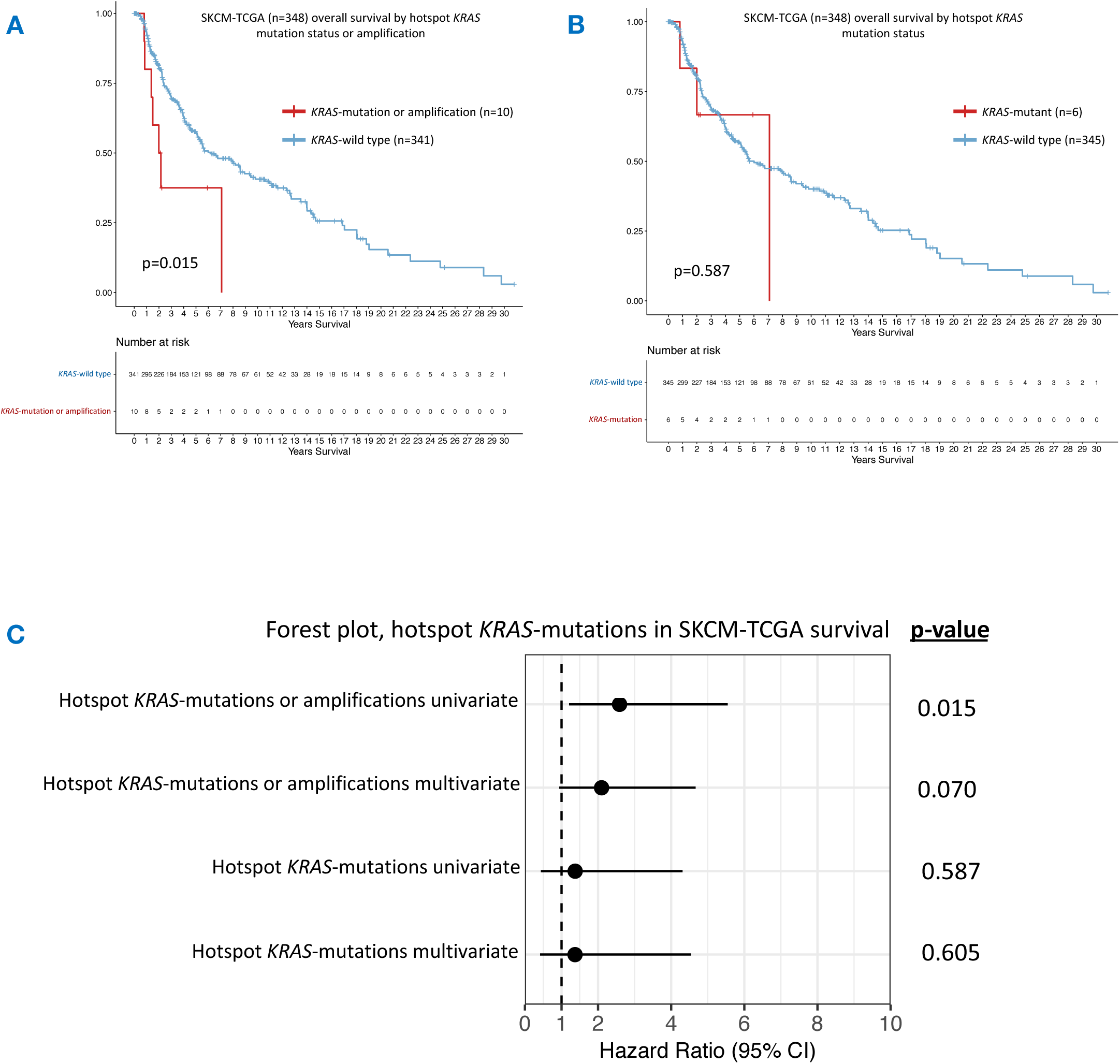
Impact of *KRAS* mutational status on overall survival in SKCM-TCGA. **A)** Kaplan-Meier curves for overall survival in *KRAS* hotpot mutations or amplifications (n=10) versus *KRAS*-WT (n=341) melanoma in SKCM-TCGA. **B)** Overall survival in *KRAS* hotpot mutation only (n=6) versus *KRAS*-WT (n=345) melanoma in SKCM-TCGA. **C)** Forest plot showing *KRAS*-mutant survival in SKCM-TCGA (either hotspot mutations or amplifications n=10, or hotpot mutations alone n=6) in univariate (HR 2.59, 95% CI 1.21-5.55, p=0.015 for mutation/amplification and HR 1.37, 95% CI 0.44-4.31, p=0.587 for mutation alone) and multivariate (HR 2.10, 95% CI 0.94-4.67, p=0.070 for mutation/amplification and HR 1.37, 95% CI 0.41-4.53, p=0.605 for mutation alone) Cox proportional hazards regression models. Multivariate correction was undertaken for stage, sex, age at diagnosis of primary, non-synonymous mutation count as well for *BRAF* and *NRAS* mutation status.

**Supplementary Table 1. Demographic and clinical characteristics of all samples in the discovery cohort (n=50).** Germline and sequenced extracranial samples (the latter with concordant *KRAS* mutations) are also indicated in column “Tissue type”.

**Supplementary Table 2. Clinical characteristics of hotspot *KRAS*-mutant (n=4) vs *KRAS*-wild type (n=46) patient within the discovery cohort.**

**Supplementary Table 3. Clinical and mutational characteristics of the patients with hotspot *KRAS* mutations in SKCM-TCGA (n=6) and SKCM-MSK-IMPACT (n=2).** *KRAS* mutations were mutually exclusive to *BRAF/NRAS/HRAS* hotspot mutations in both datasets.

**Supplementary Table 4. List of cancer driver genes (n=549) included in the custom capture bait used on the (n=18) external validation samples.**

**Supplementary Table 5. List of cancer driver genes (n=278) included in the custom capture bait used on the orthogonal validation of driver SNVs (n=60) from the discovery cohort.**

## Author contributions

R.R. coordinated the clinical and molecular data extraction, sequencing, analyzed the data and wrote the paper; P.F. coordinated the samples and clinical data extraction from both the MIA and centers across New Zealand and co-wrote the paper; K.W. performed the bioinformatic analyses including somatic variant and copy number calling and plotted the copy number profiles. C.T. and P.E. identified the appropriate cases from across New Zealand, and extracted all the samples and associated clinical data. U.M. identified the appropriate cases from New York University Medical Center and extracted all the samples and associated clinical data. J.M.S. and M.R.D. identified the appropriate cases from the University of Queensland and extracted all the samples and associated clinical data. S.R.L. (The University of Queensland), I.O. (New York University Medical Center), G.V.L. and R.A.S. (The Melanoma Institute Australia) funded and facilitated provision of patient samples, patient’s data, materials, reviewed manuscript, and provided senior input from the respective comprehensive cancer centers. I.O. was the first to describe ‘brain metastases developing as the first site of visceral relapse’^6^ and provided expert input on this phenomenon. P.C. provided clinical supervision including critical review of the manuscript. D.J.A. provided overall supervision on all aspects of the study as well as critical review of the manuscript. All authors approved the final version.

