## Supplementary Methods for "Hotspot *KRAS* mutations in brain metastases at the first metastatic recurrence of cutaneous melanoma"

**Supplemental Methods**

**Patient enrolment**

Patients with surgically resected brain metastases and no other visceral sites of metastatic disease (assessed prior to neurosurgery by CT or MRI imaging), were eligible for inclusion. Patients with available archival paraffin embedded melanoma brain metastases suitable for DNA extraction were selected from prospectively maintained databases from; The Melanoma Institute of Australia (n=34), The Wellington School of Medicine (n=8), New York University School of Medicine (n=4) and Cambridge University Hospitals (n=4) (total 50, discovery cohort). Samples from patients selected from The University of Auckland and The University of Queensland Australia (9 and 9 cases respectively) made up the external validation cohort. Two additional patients from the discovery cohort with hotspot *KRAS* mutations who had available extracranial tumor tissue for molecular analysis were also sequenced for the presence of concordant extracranial *KRAS*-mutations (patient MBM_Disc_23 sample PD42097c regional lymph node, and patient MBM_Dis_40 samples PD31211d/PD31211e both extracranial skin metastases, clinical details in **Supplementary Table 1** and sequencing data in **Supplementary Data**). All neuro-resections were undertaken between 2008 and 2018 at the respective academic neurosurgical centers as part of routine clinical care. All cases were ethically approved by the local Institutional Review Boards (REC approval reference numbers; HREC/RPAH/444, 16/CEN/149, 10362, 11/NE/0312, HREC 2005/022 and 16/CEN/149 for the six centers outlined above respectively), as well as by the Sanger Institute’s human materials and data management committee. All samples and clinical details are listed in **Supplementary Table 1**.

**Extraction of clinical details**

Patient demographics, primary tumor characteristics (date of primary diagnosis, Breslow thickness, ulceration, mitotic rate, N stage), date of diagnosis of brain metastasis, date of neurosurgical resection, sites of surgically resected brain metastases as well as date of last follow-up or death, were extracted primarily from prospectively maintained clinical research databases and, if needed, through further review of the clinical record. None of the patients received systemic therapies (included targeted and immunotherapies) prior to the resection of brain metastases.

**Extraction and quality assessment of DNA and RNA**

For each tumor, a hematoxylin and eosin-stained (H&E) section was reviewed by a consultant histopathologist (PF, RAS, CT and PE) for confirmation of the histopathological diagnosis and to identify appropriate regions for DNA extraction. All samples were obtained as either 1.0 mm diameter cores or 4um tissue sections micro-dissected from the original FFPE block. Germline DNA was extracted from micro-dissected adjacent normal tissue where available (n=36). Genomic DNA was extracted using the QIAamp FFPE Tissue kit from Qiagen according to manufacturer’s instructions.

**Whole exome-sequencing of the discovery cohort**

Exome capture was performed using the Agilent SureSelect Human All Exon V5 platform. Paired-end sequencing was performed using the Illumina HiSeq (Illumina, San Diego, CA, USA) platform at the Wellcome Sanger Institute to generate 75 bp paired-end reads. Sequencing reads were aligned using BWA-MEM (v0.7.17-r1188)^1^ to the human reference genome hs37d5. PCR duplicates, secondary read alignments, and reads that failed Illumina chastity (purity) filtering were flagged and removed prior to running variant and copy number calling. For tumour samples (n-16) where no matching germline DNA was available, a panel of 39 FFPE-extracted normals was used to filter germline variants as well as artefacts. To ensure that tumor and matched normal samples had the best reciprocal genotype match, SAMtools mpileup, followed by BCFtools gtcheck were run to detect sample concordance, potential sample swaps and contamination. The resulting median sequencing coverage in the brain metastatic samples from the discovery cohort (excluding PCR duplicates) was 48x (range 11-134x) in the tumor samples and 47x (range 6-149x) in the germline samples.

MuTect (v1.1.7)^2^ and Strelka (v2.9.2)^3^ were used to call somatic SNVs and indels, respectively. Prior to running Strelka, Manta (1.5.0)^4^ was run and candidate large indels were used as input to Strelka. The minimum base quality score for somatic and germline variant calling was set to Phred 30. The Ensembl Variant Effect Predictor^5^ was used to predict the effect of variants on genes and proteins, relative to the gene build in Ensembl version 97. To remove artefacts, MuTect variant calls were filtered using a tiered approach, such that only the SNVs that met the following criteria were reported (1) Lower coverage samples (<40x); Variant Allele Frequency (VAF) >0.1. (2) All other depth samples; requirement of at least 2 alternative bases on each strand, total depth >=5, and VAF>=0.1 Variant calls found in the gnomAD database^6^ were removed if the global population variant allele frequency was greater than or equal to 0.01. Mutational load was calculated as the number of non-synonymous mutations per Mb. The alignments for all reported variants in melanoma driver genes (**Figure 1a**) were visually inspected using JBrowse and IGV.

Of note extracranial tumor tissue was available and whole-exome sequenced for two hotpot *KRAS* mutant patients. Patient MBM_Disc_23 sample PD42097c - regional lymph node also carried a *KRAS^Q61R^* mutation, and patient MBM_Dis_40 samples PD31211d/PD31211e - extracranial skin metastases also carried *KRAS^G12D^* mutations, both concordant with the *KRAS* mutation in the matched brain metastases (raw data deposited in **Supplementary Data**).

**Copy number profiling of the discovery cohort**

Sequenza (v2.1.2)^7^ was used to estimate tumor cellularity and ploidy from 30 tumor-normal pairs in the discovery WES cohort (36/50 discovery cohort samples were paired with germline; minus 5 samples with coverage < 20x and one sample with a noisy CNV profile PD36788e), as well as to calculate allele-specific copy number profiles. For each sample, the default best-fit solution was used for the cellularity and ploidy estimates. To call copy number gain or loss in each sample, the neutral copy number was set as the weighted mean copy number of the segments, rounded to the nearest whole number.

**Targeted panel sequencing of the external validation cohort**

Panel sequencing of the 18 samples in the external validation was performed using custom pull-down and sequencing of 549 key melanoma and related cancer driver genes (**Supplementary Table 4**). A custom capture probe was designed using Agilent Technologies’ online software ‘Sure Select Design Wizard’ (ELID Design ID: 3065404). DNA capture libraries were created using native DNA (paired-end, average insert size 150bp). Libraries were multiplex sequenced using the Illumina HiSeq platform, excluding reads from PCR duplicates. Only samples with an average depth of >10x (excluding PCR duplicates) across the entire bait were included. SAMtools mpileup was used to identify non-reference bases at the *BRAF*/*NRAS*/*HRAS*/*KRAS* hotpot loci, giving a total of 44 loci for interrogation in each of the 18 (tumor-only) samples. Only non-reference variants (with minimum based quality 30 and mapping quality 10) supported by at least 2 alternate bases (from reads not marked as PCR duplicates) are reported. The median sequencing coverage (excluding PCR duplicate reads) across all these hotspot loci in the 18 external validation samples was 44x, with 700/792 (88%) of loci covered with ≥ 10 bases.

**Orthogonal validation of SNVs in the discovery cohort**

Melanoma driver single nucleotide variants (SNVs) called in the whole-exome sequenced discovery cohort were orthogonally validated (with an aliquot of the same DNA) using a custom gene panel designed to capture (n=287) cancer driver genes in solid tumors within TCGA and ICGC (**Supplementary Table 5**, ELID ID: 0822402). A variant called in the discovery cohort (through whole-exome sequencing) that was also present in the orthogonal validation custom panel (with minimum base quality 30 and mapping quality 10) and supported by at least 1 alternate bases in the validation, is reported as validated somatic. With these criteria, 56/60 (93%) of the substitutions were validated somatic (true positives) (**Supplementary Data**). The (n=4) SNVs called in the discovery cohort that were not validated in the orthogonal validation may represent subclonal mutations that require greater depth rather than false positive calls.

**Comparison of the mutational profiles to cutaneous melanomas represented in TCGA and the MSK-IMPACT datasets**

In order to compare the mutational rates and frequencies to our dataset (represented by one sample per patient), the clinical and mutation data from The Cancer Genome Atlas (SKCM-TCGA)^8^, were downloaded from the cBioPortal^9,10^. Samples were filtered to a single sample per patient giving a total of 358 samples from 358 patients, of which all samples had appropriate SNV data. The demographic characteristics of these 358 SKCM-TCGA patients were largely comparable to our cohort, whereby (n=220, 62%) were male and the median age was 57 years (95% CI 47-71 years). The majority of these samples (n=283, 79%) were classified as ‘metastasis’, as opposed to ‘primary’ (n=75, 21%). The tumor sites were mainly from skin, subcutaneous or nodal metastatic sites, including 144 (40%) classified as from ‘extremities’, 134 (37%) from ‘truncal’ locations, 26 (7%) from the ‘head and neck’, 23 (6%) were ‘regional lymph nodes’ and the remainder were from other (less frequent) metastatic sites. Only 5 tumor samples in SKCM-TCGA were classified as from the ‘brain’ and these were excluded when we subsetted to the ‘extracranial’ metastatic melanoma comparator (n=274).

The MSK-IMPACT dataset was extracted from the publication by Zehir *et al*.^11^. Samples were also filtered to a single sample per patient giving a total of 186 samples from 186 patients, all of which had appropriate SNV data. These samples were all labelled as ‘cutaneous melanomas’, and the majority (n=161, 87%) were classified as ‘metastasis’ as opposed to ‘primary’ (n=25, 13%). All primary tumors in this dataset (n=25) were classified as from ‘skin’ whereas the majority of metastatic samples were from ‘regional lymph node’ (n=25, 16%), ‘lung’ (n=24, 15%) and ‘in-transit’ (n=17, 11%) and the remainder were from other (less frequent) metastatic sites.

Both SKCM-TCGA and SKCM-MSK-IMPACT datasets included only cutaneous melanomas, in particular any melanomas within these datasets from acral, mucosal and other rarer sites were excluded. All SNVs reported from these datasets were non-synonymous mutations. We utilized the mutation calls provided by these resources and did not perform any additional filtering. The somatic mutational rate was calculated as the number of non-synonymous mutations divided by 30Mb, assuming that an average exome has 30Mb in protein-coding genes with sufficient coverage.

**Copy number profiling of the SKCM-TCGA cohort**

Copy number calls were generated from single nucleotide polymorphism (SNP data) within the SKCM-TCGA dataset using allele-specific copy number analysis (ASCAT version), as previously described.^12^

**Statistical methods**

The mutational landscape of our cohort was compared to the corresponding published landscapes using a Fisher’s exact test. All statistical analysis and graphics were generated using R version 3.5.1 (R Foundation for Statistical Computing, Vienna, Austria. URL [http://www.R-project.org/](http://www.r-project.org/)).

**Survival analyses**

Overall survival from resection of brain metastasis was calculated as the time from the date of resection of the brain metastasis to last follow-up (censored) or death from any cause. Overall survival from primary tumor was calculated as the time from the date of resection of the primary tumor to last follow-up (censored) or death from any cause. Univariate Kaplan-Meier survival curves were constructed using the R ‘Survival’ package version 2.4.2. Univariate and multivariate hazard ratios (HR), 95% confidence intervals (95% CI) and corresponding p-values were obtained using Cox proportional hazards regression models. Multivariate correction was undertaken for gender, age, as well as *BRAF* and *NRAS* mutation status.

**Supplementary data**

All the whole exome and targeted sequencing data (including raw sequencing files, variant calls and copy number calls) have been deposited at the European Genome-Phenome Archive (<https://www.ebi.ac.uk/ega/> at the EBI).

**Supplemental methods references**

1.	Li H. Aligning sequence reads, clone sequences and assembly contigs with BWA-MEM. *arXiv.* 2013;1303.
